# Phosphine: an eco-friendly alternative for management of wheat storage insects

**DOI:** 10.1101/2020.03.05.978486

**Authors:** Sumitra Arora, J. Stanley, C. Srivastava

## Abstract

Methyl bromide (MB) fumigant has been phased out in many countries honouring ‘Montreal Protocol’. Reports on phosphine fumigant efficacy under different ecological zones in India are scanty. Field fumigation trials were conducted on 5 Metric Tons of wheat stacks per replication of a treatment for testing phosphine efficacy against laboratory cultured and resident population of *Tribolium castanium*, *Sitophilus oryzae*, and *Rhyzopertha dominica*. The trials were organized in triplicate including control at two locations with varying climate locations namely, Pithoragarh and Delhi using conventional 56% tablet (2 and 3 tablet/MT) and 77.5% granular (1.0 and 1.5 g phosphine/m^3^) formulations of Aluminium phosphide (AlP) for 7 and 10-days exposure periods for each dosage. Phosphine concentration was monitored every 24 hours till end of the exposure periods. Insect mortality of laboratory culture and resident infestation was observed as 100% in both the exposure periods using all dosages of phosphine. No emergence of insects was observed till 60 days after treatment at both the locations, indicating mortality of all available life stages of insects during the exposure periods. Minimum effective phosphine concentration for controlling all life stages of *S. oryzae*, *R. dominica* and *T castaneum* was observed 500-600 ppm for 7-days exposure at 25°-27°C and humidity 38-45%, at 1.5 g phosphine/m^3^. Hence, phosphine can be an effective alternative to methyl bromide fumigant for wheat stored grain insects at different climatic conditions.

## 1. Introduction

Stored food grain insects are serious problems of dried, stored, durable agricultural commodities and of many value-added food products and non-food derivatives of agricultural products worldwide which can cause serious post-harvest losses, estimated up to 9 % in developed countries to 20 % or more in developing countries (Pimentel, 1991). Wheat is an important staple food of India and other Asian countries. India has an estimate production of wheat about 102 MT for 2018-19 (DAC, 2018). The produce is stored for 2-12 months by public sector agencies like Food Corporation of India, the Central and State Warehousing Corporation, before distribution through Public Distribution System. Wheat is stored in jute bags of 50 kg capacity in these warehouses.

During storage, wheat grains are attacked by a number of storage insect pests, such as, *Tribolium castanium* (red floor beetle, adults and larvae)*, Sitophilus oryzae* (grain weevil), and *Rhyzopertha dominica* (lesser grain borer), responsible for post-harvest losses. Food and Agriculture Organization of U.N. predicts about 1.3 billion tons of food are globally wasted or lost per year (Gustavasson, et al. 2011). Out of a total 10% post-harvest losses of grains, a significant 6% are damaged during their storage. Fumigation is the primary option for controlling these insects. Methyl bromide (MB) had been used since years as an effective broad spectrum fumigant. However, phasing out of MB globally honouring ‘Montreal Protocol’ due to its ozone depletion properties, led to the exploration of other alternative fumigants.

Phosphine is used worldwide and is a common fumigant registered and used in India as aluminium phosphide formulations for domestic storage, and needs testing for long term storage and Quarantine shipments. In India, conventional 56% Aluminium phosphide (AlP) is available as 3g tablet, and 10g as well as 34g pouches; and 77.5% as granules. The 6% tablet is recommended for treatment in burrows for field rodents. Another source of phosphine is Magnesium phosphide plates which has been registered for fumigation of unmanufactured tobacco for export as per importing country requirement. Though there are many studies on the efficacy of phosphine against major storage pests all around the world under laboratory conditions, studies on efficacy of phosphine in warehouse storage under varying climatic conditions is limited.

Present study has been investigated for the efficacy of phosphine against stored wheat grain insects with improved formulations, so that it can be used for Quarantine Purpose and subsequently for long term storage in India. The efficacy of AlP was tested against laboratory cultured insect population of *Tribolium castanium*, *Sitophilus oryzae*, and *Rhyzopertha dominica* as well as their field populations.

## 2. Methods

### 2.1 Experimental layout

Under this study of alternatives to methyl bromide fumigant, phosphine was tested for its efficacy against stored wheat grain insect at two locations, namely New Delhi and Pithoragarh, Uttrakhand, with varied climatic conditions. Accordingly, Delhi has extreme summers and winters, and Pithoragarh has temperate climatic conditions. The on-site gas generation (through QuickPHlo-R^TM^ gas generator) using granular 77.5% formulation at 1.0 and 1.5 g phosphine/m^3^ has been used for fumigation of wheat at both the locations. The same was compared with the conventional 56% AlP tablet at 2 Tab/MT (~1.5 g phosphine/m^3^) and 3 Tab/MT (~2.3 g phosphine/m^3^). The experiments were conducted on 150 MT of wheat grains, piled in 30 stacks for 7 and 10-days exposure periods during 2017-18 (stacks in triplicate for each dosage and exposure period including control). The stacks of wheat grains, each of 5 MT, comprised of 100 bags (50 kg each) of grains. Three layers of monitoring lines were positioned in each stack including controls, for recording phosphine gas concentration during the experiment.

Temperature and relative humidity were recorded every 24 hours during the course of field trials at each location. Moisture content of the wheat grains was also determined before initiating the field fumigation trials.

### 2.2 Rearing of Test insects

The rust-red flour beetle, *Tribolium castaneum* was reared in whole wheat flour. The rice weevil, *Sitophilus oryzae* and the lesser grain borer, *Rhyzopertha dominica* were cultured in wheat grains kept in big plastic containers. The grains were washed, sun dried and infestations, if any were removed, before culturing the insects. Pure culture of rice weevil and lesser grain borer were kept in containers and reared separately. The culture was provided with fresh food grains as and when required.

### 2.3 Field fumigation trials of wheat stacks and gas concentration monitoring

To evaluate the efficacy of treatments, tests insects enclosed in plastic containers with perforated lids were used. Test insects included 10 each of available life stages of *T. castanium*, *S. oryzae*, and *R. dominica* in 12 test containers placed uniformly in three levels (top, middle and bottom) across four sides of the stacks.

The wheat grain stacks were fumigated with 77.5% granular formulation at two dosages, 1.0 and 1.5 g phosphine/m^3^, using on site QuickPHlo-R™ phosphine generator (United Phosphorus Ltd., India); and conventional 56% AlP tablet formulations at 1.5 g phosphine/m^3^ (2 tab/MT) and 2.3 g phosphine/m^3^ (3 tab/MT); each in triplicate including control, with no treatment. The experiments were conducted for 7 and 10-days exposure periods for both the formulations. Three replicates/stacks were designated for each dosage and exposure period including the untreated control.

The conventional AlP tablet formulation (both dosages) was wrapped in muslin cloth pouch before positioning them uniformly in three layers of each stack. The stacks were covered with fumigation cover followed by sealing using sand bags. For granular 77.5% formulation, stacks were sealed prior to on-site gas generation, with complete release of phosphine gas in 60-minute cycle. Leakage of phosphine gas was checked/detected using Phosphine Alert^®^ (UPL; with measuring range 0.2-20 ppm), and rectified wherever required. Phosphine gas concentration was monitored every 24 hours throughout the experimental period, using UNIPHOS FumiSense Pro (UPL; measuring range 1-2000 ppm) through gas monitoring lines. For on-site gas generation, concentration monitoring was initiated on zero day itself one hour after fumigation; while, in case of conventional tablet, it was initiated one day after placement of tablets i.e, treatment.

### 2.4 Efficacy of phosphine against test insects

On termination of fumigation after each exposure period, test insect containers were retrieved from the stacks and monitored for insect survival/emergence in laboratory. Representative grain samples (250 g) were also drawn from each stack (including control) before start and on termination of the treatments for infestation monitoring of laboratory as well as field population. Observations were made immediately (within a week interval), 30 and 60-days post treatments.

## 3. Results

Average relative humidity and temperature at Pithoragarh, measured during the field fumigation trials, was 33-43 % and 22-26°C respectively, during 1^st^ week of October, 2017; while at Delhi it was 40-50 % and 25-27°C respectively, during mid-week of November, 2017. The moisture in wheat grains was monitored before the fumigation trials and average moisture content in wheat grains was 12.5 and 10.5% before fumigation at Pithoragarh and Delhi, respectively.

### 3.1 Gas concentration monitoring

The concentration monitored during the fumigation exposure periods regularly for 7 and 10 days are presented in Table 1. At Pithoragarh site, the terminal phosphine gas concentration observed for granular 77.5% formulations at 1.0 and 1.5 g phosphine/m^3^, were 344 and 499 ppm for 7-days; and 210 and 181 ppm for 10-days exposure periods, respectively. For conventional 56% AlP tablet formulation, the terminal concentrations at 1.5 (2 tab/MT) and 2.3 (3 tab/MT) g phosphine/m^3^ were 547 and 990 ppm for 7-days; and 507 and 434 ppm for 10-days exposure periods, respectively. However, at Delhi site, the terminal concentrations of phosphine were 328 and 229 for 77.5% G formulation at 1.0 g phosphine/m^3^; and 640 and 424 at 1.5 g phosphine/m^3^ after 7 and 10 days of treatment, respectively. The concentrations of phosphine were 555 and 478 ppm with 56% AlP tablet at 1.54 g/m^3^; and 1163 and 749 ppm at 2.3 g/m^3^, after 7 and 10 days of treatment, respectively (Table 1). The peak phosphine concentrations were observed on 2^nd^ / 3^rd^ day of AlP 56% tablet treatments.

**Table 1.** Phosphine gas concentration profile in wheat stacks fumigated with AlP formulations

| Location | Days after fumigation | Phosphine concentrations (ppm)* (±SE) |  |  |  |  |  |  |  |
| --- | --- | --- | --- | --- | --- | --- | --- | --- | --- |
|  |  | 7-day exposure<sup>#</sup> |  |  |  | 10-days exposure<sup>#</sup> |  |  |  |
|  |  | Tablet application |  | On site generation |  | Tablet application |  | On site generation |  |
|  |  | 1.54 g/m<sup>3</sup><br>(2 tab/MT) | 2.31 g/m<sup>3</sup><br>(3 tab/MT) | 1.0 g/m<sup>3</sup> | 1.5 g/m<sup>3</sup> | 1.54 g/m<sup>3</sup><br>(2 tab/MT) | 2.31 g/m<sup>3</sup><br>(3 tab/MT) | 1.0 g/m<sup>3</sup> | 1.5 g/m<sup>3</sup> |
| Pithoragarh | 0 | - | - | 1280±34.6 (103.8) | 1690±56.3 (168.9) | - | - | 1320±38.5 (115.5) | 1710±47.1 (141.3) |
|  | 1 | 410±12.3 (36.8) | 718±48.9 (146.8) | 1057±26.3 (69.6) | 1468±48.5 (168.0) | 316±34.4 (84.4) | 600±12.4 (35.0) | 1142±36.0 (108.1) | 1406±13.1 (39.3) |
|  | 2 | 943±39.6 (118.6) | 1374±45.3 (135.8) | 866±27.4 (82.1) | 1186±37.3 (111.9) | 927±24.6 (65.1) | 1287±12.7 (35.8) | 860±13.9 (41.7) | 1096±13.2 (39.9) |
|  | 3 | 1082±55.1 (165.2) | 1647±43.4 (130.1) | 727±14.4 (40.7) | 1053±27.0 (85.5) | 1074±23.8 (67.3) | 1533±8.9 (23.6) | 678±22.1 (62.5) | 919±15.7 (44.7) |
|  | 4 | 836±61.3 (183.8) | 1627±24.9 (74.8) | 602±21.4 (64.2) | 885±22.5 (71.2) | 959±22.2 (58.8) | 1503±27.5 (72.8) | 586±15.2 (45.6) | 763±5.4 (13.2) |
|  | 5 | 817±53.7<br>(151.8) | 1349±50.0<br>(150.1) | 422±41.7<br>(102.1) | 730±21.7<br>(68.7) | 898±16.0<br>(48.1) | 1289±38.7<br>(102.3) | 473±22.5<br>(67.6) | 650±15.1<br>(39.8) |
|  | 6 | 669±38.9<br>(110.1) | 1125±34.4<br>(103.3) | 397±24.3<br>(72.8) | 661±9.2<br>(27.7) | 772±13.6<br>(38.5) | 1055±26.4<br>(79.1) | 433±12.0<br>(36.1) | 584±9.8<br>(26.1) |
|  | 7 | 547±27.9<br>(73.9) | 990±36.6<br>(109.9) | 344±27.4<br>(72.4) | 499±13.9<br>(44.2) | 677±10.5<br>(27.7) | 857±22.2<br>(62.7) | 356±10.3<br>(31.0) | 402±13.7<br>(38.9) |
|  | 8 |  |  |  |  | 612±11.0<br>(33.1) | 691±9.3<br>(24.5) | 282±9.3<br>(27.9) | 319±16.6<br>(47.1) |
|  | 9 |  |  |  |  | 535±25.3<br>(75.9) | 563±23.4<br>(70.1) | 237±9.7<br>(29.0) | 230±1.2<br>(3.7) |
|  | 10 |  |  |  |  | 507±31.2<br>(93.5) | 434±22.2<br>(66.6) | 210±3.3<br>(9.8) | 181±3.2<br>(10.3) |
| Delhi | 0 | - | - | 930±13.9<br>(41.7) | 1581±39.6<br>(118.9) | - | - | 938±15.8<br>(47.3) | 1689±41.1<br>(123.3) |
|  | 1 | 798±11.3<br>(33.9) | 901±11.3<br>(33.9) | 732±18.8<br>(56.3) | 1338±27.9<br>(83.8) | 702±22.6<br>(67.7) | 978±32.1<br>(96.2) | 729±19.6<br>(58.9) | 1397±14.2<br>(42.6) |
|  | 2 | 1089±26.9<br>(80.6) | 1478±33.5<br>(100.5) | 672±10.1<br>(30.2) | 1205±17.8<br>(53.5) | 1065±34.5<br>(103.5) | 1556±48.1<br>(144.4) | 630±16.1<br>(48.4) | 1237±13.9<br>(41.8) |
|  | 3 | 1109±49.6<br>(148.8) | 1687±45.3<br>(136.0) | 596±13.5<br>(40.6) | 1061±18.9<br>(56.8) | 1052±36.0<br>(108.1) | 1646±56.2<br>(168.6) | 560±6.9<br>(20.8) | 1071±15.0<br>(44.9) |
|  | 4 | 840±21.7<br>(65.2) | 1548±39.7<br>(119.0) | 518±20.1<br>(60.2) | 984±15.7<br>(47.0) | 998±19.4<br>(58.2) | 1609±72.0<br>(216.0) | 474±7.2<br>(21.7) | 946±11.0<br>(33.1) |
|  | 5 | 782±64.0<br>(192.1) | 1489±68.9<br>(206.6) | 467±21.0<br>(63.0) | 830±16.1<br>(48.3) | 861±11.9<br>(35.6) | 1428±34.6<br>(103.8) | 428±19.8<br>(59.3) | 822±13.9<br>(41.8) |
|  | 6 | 658±47.2<br>(141.6) | 1313±79.4<br>(238.3) | 398±19.0<br>(57.0) | 729±16.1<br>(48.3) | 703±19.7<br>(59.1) | 1246±67.9<br>(203.7) | 353±6.1<br>(18.4) | 713±13.5<br>(40.6) |
|  | 7 | 555±10.4<br>(31.2) | 1163±24.2<br>(72.5) | 328±18.6<br>(55.9) | 640±14.3<br>(42.9) | 628±16.6<br>(49.8) | 1092±64.7<br>(194.1) | 301±5.2<br>(15.5) | 636±12.4<br>(37.1) |
|  | 8 |  |  |  |  | 557±13.8<br>(41.4) | 974±36.2<br>(108.6) | 263±5.5<br>(16.4) | 563±11.5<br>(34.4) |
|  | 9 |  |  |  |  | 478±13.4<br>(40.2) | 851±32.9<br>(98.8) | 229±4.5<br>(13.5) | 491±10.7<br>(32.0) |
|  | 10 |  |  |  |  | 414±14.8<br>(44.4) | 749±29.9<br>(89.6) | 198±4.0<br>(11.9) | 424±10.0<br>(29.9) |

\*Average of 3 stacks as measured with FumiSense Pro
#temperature range: 22-27°C; RH: 33-43% at Pithoragarh; and
#temperature range: 25-27°C; RH: 40-50% during the experiment trials

### 3.2 Efficacy of phosphine against storage insects

The efficacy of AlP was evaluated against laboratory cultured insect population of *T. castaneum*, *S. oryzae*, and *R. dominica* as well as the field/resident populations.

#### 3.2.1 Insect mortality of resident population in fumigated stacks

The insect mortality in wheat grain samples drawn before fumigation was compared with those of insects in post fumigated grain samples after 7, 30 and 60 days of fumigation. As per records of the warehouse, the wheat stacks at Pithoragarh received from Haryana on 16 June 2017, were already fumigated; and followed by another fumigation at FCI Pithoragarh during last week of August, 2017, before initiating our field trials. Hence, no insect population was found in treated or untreated grain samples under observations recorded on 7, 30 and 60 days after treatments. Only few (1-2 in number) live *S. oryzae* insects were found in wheat samples drawn before fumigation. Contrarily, the wheat stacks at Delhi were heavily infested with adults and larvae population of *T. castaneum*. However, 100% insect mortality was recorded for resident population in fumigated wheat stacks at Delhi. Besides this, insect count in untreated control samples of wheat grains, and samples drawn before fumigation was also recorded and analysed statistically (Table 2 and 3).

**Table 2.**
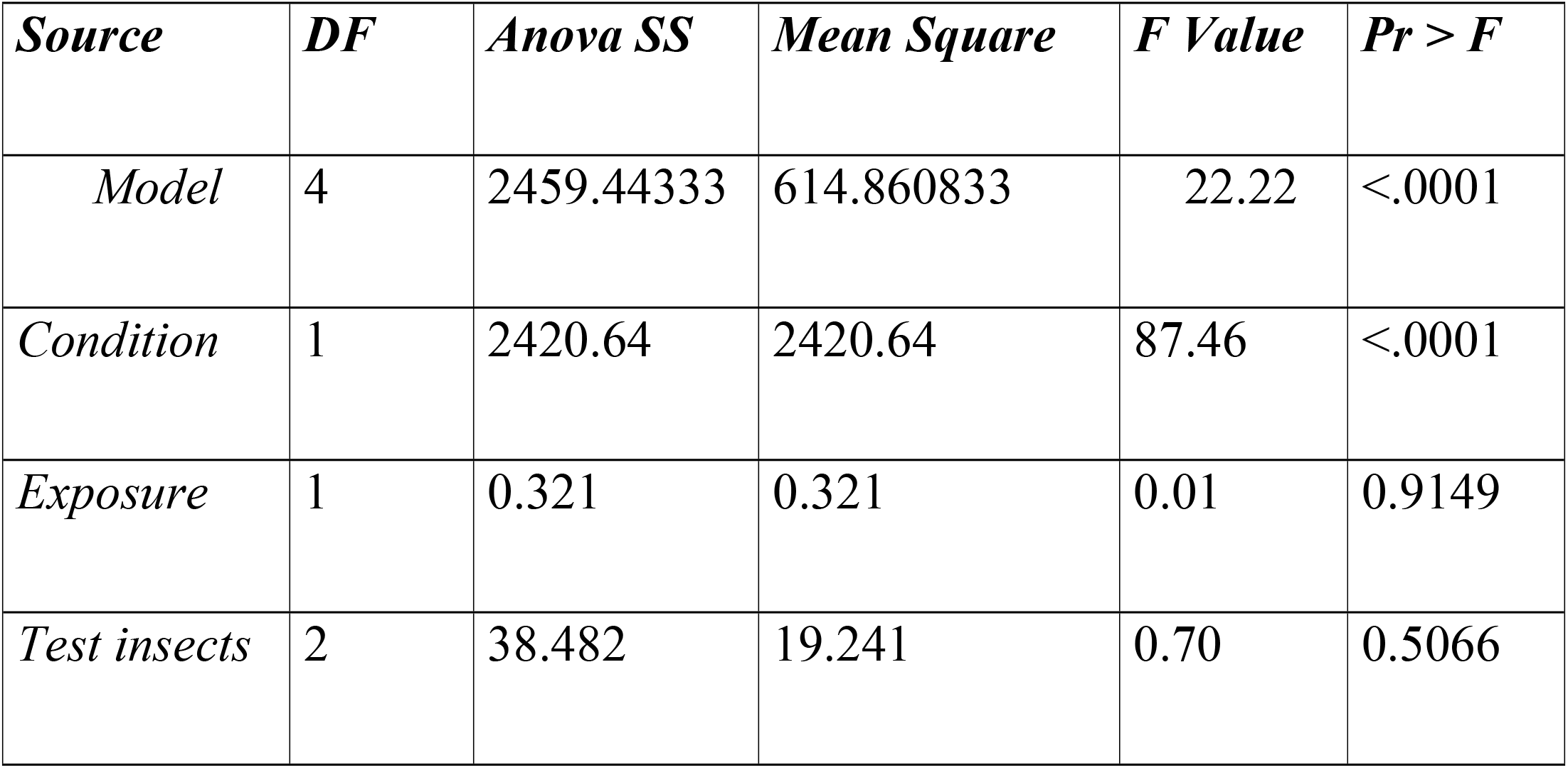
ANOVA for insect count in Pre (before fumigation) and control (no treatment) grain samples at two exposure levels

| *Source* | *DF* | *Anova SS* | *Mean Square* | *F Value* | *Pr > F* |
| --- | --- | --- | --- | --- | --- |
| *Model* | 4 | 2459.44333 | 614.860833 | 22.22 | <.0001 |
| *Condition* | 1 | 2420.64 | 2420.64 | 87.46 | <.0001 |
| *Exposure* | 1 | 0.321 | 0.321 | 0.01 | 0.9149 |
| *Test insects* | 2 | 38.482 | 19.241 | 0.70 | 0.5066 |

**Table 3.**
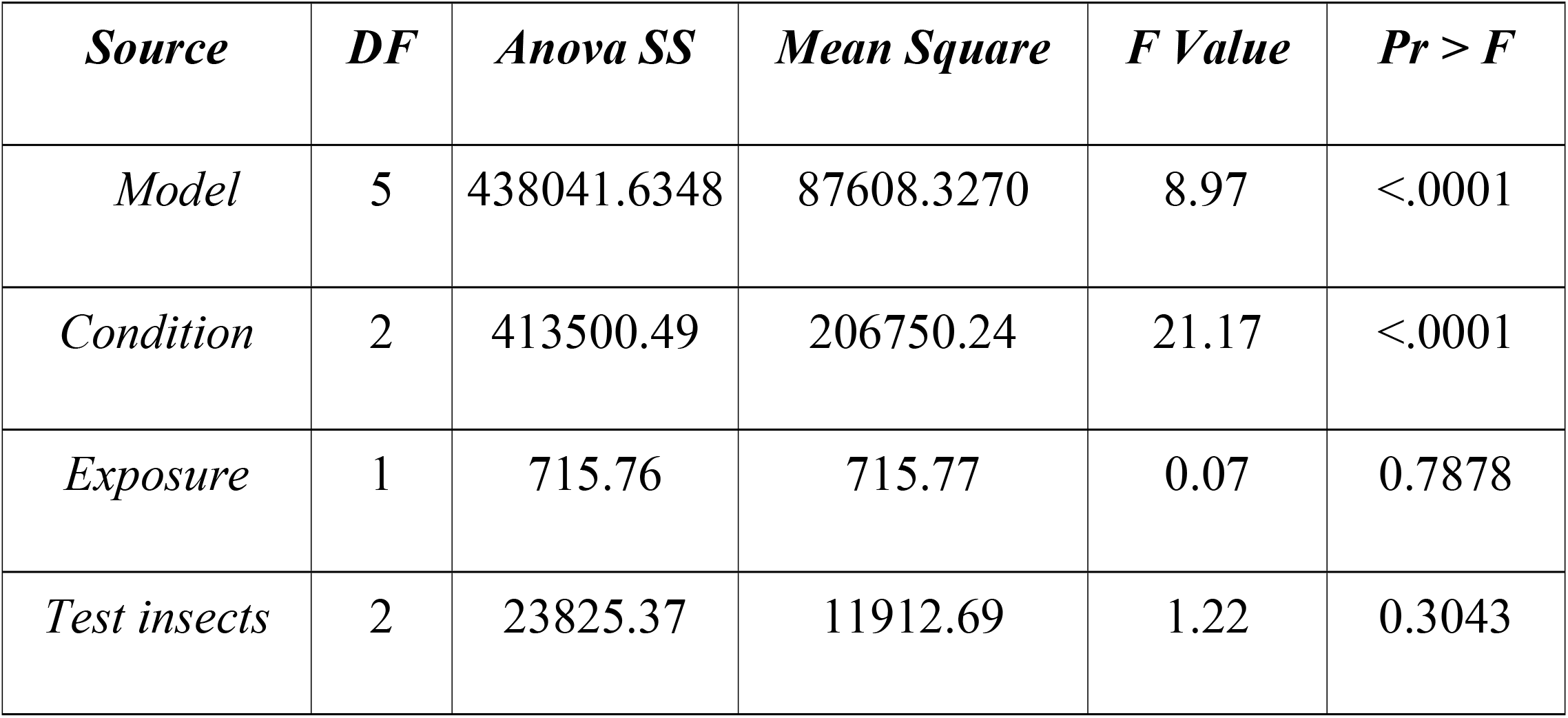
ANOVA for comparison of test insects at two locations

| *Source* | *DF* | *Anova SS* | *Mean Square* | *F Value* | *Pr > F* |
| --- | --- | --- | --- | --- | --- |
| *Model* | 5 | 438041.6348 | 87608.3270 | 8.97 | <.0001 |
| *Condition* | 2 | 413500.49 | 206750.24 | 21.17 | <.0001 |
| *Exposure* | 1 | 715.76 | 715.77 | 0.07 | 0.7878 |
| *Test insects* | 2 | 23825.37 | 11912.69 | 1.22 | 0.3043 |

#### 3.2.2 Monitoring of laboratory culture insect population placed in wheat grain stacks

The test containers with laboratory cultured insects, placed in stacks for fumigation, were taken out for insect survival/emergence count after respective exposure periods. The laboratory cultured insects were monitored for survival or emergence of next generation adults and larval stages of *T. castaneum,* adults of *S. oryzae,* and *R. dominica* in post treated and untreated (control) grain samples. All available stages of insects in test containers were found dead in all the treatments, except control, at both locations, Insect mortality in control treatments was observed in the range of 5-25% with *S. oryzae* adults and *T. castaneum* larvae. The laboratory cultured live insect populations were recorded in untreated control samples of wheat grains, and analysed statistically using software SAS (version 9.4) (Table 2 and 3).

## 4. Discussion

For effective fumigation treatment the field variables such as temperature, gas concentration, insect tolerance and grain types are taken into consideration for dosage regimens using any fumigant (EPPO, 1984; AFHB/ACIAR, 1989). The major impediments for the existing fumigation technology is phasing out of methyl bromide due to its ozone depletion properties. Research priorities are, hence, focussing on the development of methods to minimize atmospheric emission of methyl bromide and to find out its suitable substitutes. Phosphine is the most widely used and reported as an effective fumigant in the world. It is the preferred chemical for routine grain disinfestations in the developing countries where other alternative techniques, such as controlled atmosphere storage, are expensive or cannot be readily adopted. Phosphine has many desirable properties of a fumigant with a few disadvantages which could be minimised by maintaining low moisture content of the grains to be fumigated and practicing proper aeration after fumigation (Rajendran and Gunasekaran, 1995).

### 4.1 Phosphine gas concentration versus climatic conditions

The peak concentration in case of conventional 56% AlP tablet, was observed after 3 days (72 hours) of fumigation, due to very low humidity (33-43%) as well as low temperature (22-26°C) at Pithoragarh location. However, the gas release, through on site gas generation using granular 77.5% AlP, is not affected where complete gas is released within 60 minutes of fumigation using gas generator. The humidity at Delhi centre was little higher than Pithoragarh during the field experimentation trials; however, not enough to effect the phosphine release. The peak concentration in case of conventional 56% tablet, was observed after 3 days (72 hours) of fumigation, due to low humidity (40-50%) as well as low temperature (25-29°C) at Delhi location. The onsite gas generation using granular 77.5% AlP released peak concentration on zero day itself.

The terminal concentrations for conventional tablet and on-site gas generation cannot be similar at both the locations, as the peak concentration for on-site gas generation is achieved on zero day itself; while it is dependent on temperature and humidity in case of release of phosphine from tablet formulation. The minimum effective phosphine concentration for controlling all life stages of *S. oryzae*, *R. dominica* and *T castaneum* was observed as 500-550 ppm for 7-days exposure at 25°C temperature and average RH 38%, using conventional tablet or on-site gas generation at 1.5 g phosphine/m^3^ at Pithoragarh site; while it was observed slightly higher, 550-600 ppm, at 27°C and humidity 45% using both the phosphine formulations with same dosage at Delhi location. No significant difference in terminal phosphine concentration was observed for 1.5 g phosphine/m^3^ with 7 days exposure period in both the locations. Winks and Hyne (1997) reported minimum effective phosphine concentration for susceptible strains of *Sitophilus* sp. as 700 ppm and for *R. dominica* and *T castaneum* as 300 ppm in a 7 day treatment at 20°C. However, 720 ppm of gas concentration was observed as relatively high for resistant *S. oryzae* but inadequate for controlling *R. dominica* (Rajendran and Gunasekran, 2002). The peak concentration using conventional tablet formulation achieved on 3^rd^ day is at par with the concentration at zero day with on-site gas generation using granular 77.5% AlP. However, the end concentration of phosphine after 7 and 10 days’ exposure period was sufficient to kill all available life stages of storage insects, as per recorded observation under field trials at both the locations.

Grain moisture is very much affected by climatic conditions. In present study, it was higher in wheat grains at Pithoragarh than that of Delhi location, because of relatively lower temperature and humidity. Environmental temperature and relative humidity are two most important and interdependent factors subsequently effecting the grain moisture (Strelec et al., 2010; Harrington, 1973). Harrington reported three “rules of thumbs” regarding optimal seed storage. The first one for the seeds with grain moisture from 5 to 14 % states that each 1 % reduction of seed moisture doubles the storage life of the seeds; the second one states that for each 10 °F (5.6 °C) decrease in seed storage temperature storage life of seed is doubled; and the third one states that arithmetic sum of relative humidity and storage temperature should not exceed 100 for safe seed storage, or 120 as later reported (Bewley and Black, 1985; Copeland and McDonald, 1999; Harrington, 1973). Temperatures below 21°C and relative humidity values no higher than 60% or 70% for seeds kept at 4°C should be used in implementation of these rules for long-term storage of wheat seeds, from 3 to 10 years. Wheat stored in normal warehousing conditions faces fluctuation in temperatures and humidity depending on the seasons. Under such conditions grains kept in bags easily exchange moisture with the surrounding air of defined relative humidity and temperature depending on species (Copeland and McDonald, 1999; Volenik et al., 2006). Reports have documented that if moisture equilibration between seeds and environmental air result in an increase in seed moisture, deteriorative process increases with concomitant temperature within seeds speeds up, consequently leading to germination and vigour decrease. Strelec et al. (2010) indicated significant influence of storage conditions on moisture content, germination and vigour changes during storage of wheat seeds, as well as varietal dependence of seed viability.

### 4.2 Phosphine efficacy against test insects

The emergence and survival of insects was monitored in wheat grain samples of untreated control, before (pre-) and after (post-) fumigation for 7, 30 and 60 days under simulated laboratory conditions. The laboratory cultured population of test insects in post fumigated samples of wheat grains at Delhi and Pithoragarh location was observed with 100% mortality for 7- and 10-days exposure periods, with all fumigated dosages using conventional 56% AlP tablet and 77.5% granular formulations. Similar was the case with resident population of insects in wheat grain samples at both the locations. The emergence of stored grain insects was observed in control (untreated) as well as in the samples of wheat grains taken from the stacks before fumigation at both the locations. However, no emergence/survival of insects was observed in any of the post fumigated samples after 30 and 60 days of treatment. Average mortality of insects in grain samples of untreated control stacks ranged 5-25%. Insect count of *S. oryzae* and *R. dominica* emerged in untreated control samples of wheat at Pithoragarh location was more (> 400, ~350, respectively in number) than that of Delhi (< 350, ~200 respectively in number). Reason may be the grain moisture less than 12%, which might have inhibited growth of these insects.

Analysis of variance was performed for experimental results obtained with two conditions (Pre-fumigation and control separately) and at two exposure levels (7 Days and 10 Days) and with three test insects *viz.*, *T. castaneum, R. dominica and S. oryzae.* The ANOVA (Table 2) shows a significant variation with respect to conditions and test insects, i.e. Pr > F. While the F value for the exposure levels are lower than the tabular values showing no significant variation between the exposure levels, i.e, 7 and 10 days with individual mean values being 74.506 (10 Days) and 74.317 (7 Days) and the Least Significant Difference (LSD) is 3.576.

ANOVA (Table 3) for comparison of test insect’s population at Delhi and Pithoragarh locations, and the laboratory cultured test insects, shows no significant variation in the exposure periods *viz*., 10 and 7 Days (10 days mean: 185.37; 7 days mean: 192.66; LSD: 54.08). In case of the test insects, all the three insects did not behave significantly different to each other with the LSD value 66.23. Critical value of t as 2.01.

However, under the three conditions i.e, insect population at Delhi, insect population at Pithoragarh, and Laboratory cultured insect population behaved significantly different with mean values of laboratory population = 263.67, Pithoragarh population*=* 237.17 and Delhi population = 66.21 and LSD value of 66.23. The box plot (Fig 1) clearly shows the difference in the insect populations.

**Fig. 1.**
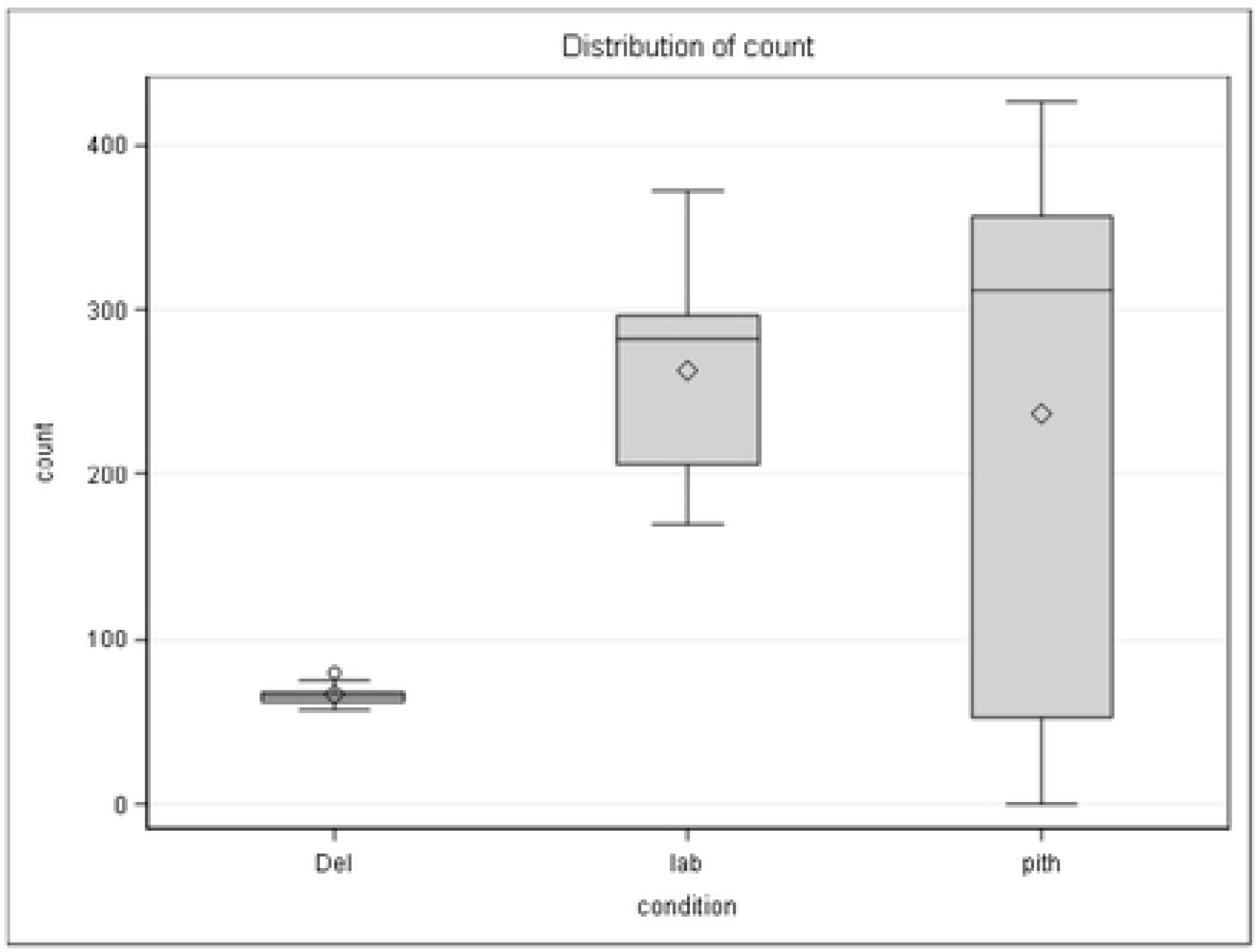
Variation in insect populations under three conditions (Del: Delhi. Pith: Pithoragarh; lab: laboratory; condition: condition (populations)

As per FAO (1994) most storage insects are able to survive and multiply rapidly on well-dried grain. However, grain dried to below 12% moisture content (mc) inhibits the development of most species to some extent and on exceptionally dry grain (<8% mc) the grain weevils, for example, are insignificant pests. The grain borers remain of considerable importance at these low moisture levels. However, population of *T. castaneum* at Delhi location kept increasing up to 60 days of post fumigation (> 350 in number) as compared to Pithoragarh location, which continued to decrease (~ 350 to 75 in number). This may be due to the climatic conditions on the survival of the insects, as Pithoragarh experienced very low temperature during that time. Temperature has been reported as the major factor for affecting insect development and population growth rates (FAO, 1994). Grain borers have been observed to be more resistant than grain weevils, but temperatures above 45°C are eventually fatal to all storage insects.

## 5. Conclusion

Release of phosphine gas is correlated with temperature and relative humidity of the environment. Grain moisture is also affected by climatic conditions. Based on the wheat fumigation trials at two locations phosphine, applied in bulk, has been observed as an effective alternative to methyl bromide fumigant in quarantine and QPS of wheat grains for controlling stored grain insects. However, maintenance of effective gas concentrations (ppm) throughout the fumigation period is essential for successful phosphine fumigation. The condition of the fumigation cover, sealing standards and the type of floor surface determine the phosphine levels and thereby insect control in the stacks.

Gas monitoring is important during fumigation to predict its success and to supplement dosage or to extend the exposure period, if necessary. For phosphine fumigations the success is ascertained based on the final-day/terminal concentration of 100 ppm and higher. Residues are formed in fumigated grains and the residue levels should not exceed the national and international tolerances.

## Acknowledgements

This work was supported and funded by Department of Agriculture and Cooperation (DAC), Ministry of Agriculture and Farmers Welfare, Govt. of India (Grant No. 8-35/2017-PP-II).

Authors are thankful to DAC, Ministry of Agriculture and Farmers Welfare, Govt of India for funding this project; and Dr PK Chakrabarty, ASRB, Member and Ex-ADG (PP and BS), ICAR for facilitating the research activities under this project. We are also thankful to Mr. Ujjwal Kumar, Business Head and his team, UPL India Pvt Ltd for helping in arranging the infrastructure, and Dr S. Rajendran, Ex-Scientist, CSIR-CFTRI and Mr IC Chadhha, Ex-Manager, CWC, for guidance and technical help as consultants. Authors also appreciate the help and cooperation received from FCI team at Delhi and Pithoragarh locations. Project team is also thankful to Directors of ICAR-NCIPM and ICAR-VPKAS. Dr. Susheel Sarkar, Scientist, ICAR-IASRI, also deserve appreciation in helping statistical analysis of the data.

